# Learning and memory deficits produced by aspartame are heritable via the paternal lineage

**DOI:** 10.1101/2023.07.09.548248

**Authors:** Sara K. Jones, Deirdre M. McCarthy, Gregg D. Stanwood, Chris Schatschneider, Pradeep G. Bhide

## Abstract

Environmental exposures produce heritable traits that can linger in the population for one or two generations. Millions of individuals consume substances such as artificial sweeteners daily that are declared safe by regulatory agencies without evaluation of their potential heritable effects. We show that consumption of aspartame, an FDA-approved artificial sweetener, daily for up to 16-weeks at doses equivalent to only 7-15% of the FDA recommended maximum daily intake value (equivalent to 2-4 small, 8 oz diet soda drinks per day) produces significant spatial learning and memory deficits in mice. Moreover, the cognitive deficits are transmitted to male and female descendants along the paternal lineage suggesting that aspartame’s adverse cognitive effects are heritable, and that they are more pervasive than current estimates, which consider effects in the directly exposed individuals only. Traditionally, deleterious environmental exposures of pregnant and nursing women are viewed as risk factors for the health of future generations. Environmental exposures of men are not considered to pose similar risks. Our findings suggest that environmental exposures of men can produce adverse impact on mental health of future generations and demonstrate the need for considering heritable effects via the paternal lineage as part of the regulatory evaluations of artificial sweeteners.

## Introduction

Emerging evidence from multiple fields of biology demonstrates that the impact of environmental exposures may extend well beyond the directly exposed individuals and manifest in multiple generations descending from the directly exposed individuals ^1–5^. Much of the research on this topic is focused on deleterious effects produced by exposure to psychosocial stress, natural calamities such as famine, or to substances such as endocrine disrupting agents, hormones and drugs of abuse ^1–3, 6–22^. However, millions of individuals are exposed daily to substances declared “safe” by regulatory agencies based on evaluation of effects in individuals directly exposed to the substances as adults or exposed prenatally.

Artificial sweeteners are one example of substances declared “safe” by regulatory agencies without a formal evaluation of potential heritable effects. Aspartame, one of the most widely consumed artificial sweeteners was approved by the US Food and Drug Administration (FDA) for use in tabletop sweeteners in 1981, carbonated drinks in 1983 and in other food products in 1996 ^23–25^. The approval appears to be based on lack of conclusive evidence for aspartame’s adverse effects in the directly exposed individuals ^23, 25–28^. However, the recent World Health Organization guidelines point out associations between consumption of aspartame and other artificial sweeteners and increased risk for metabolic disease, cardiovascular disease and cancer ^29^. Interestingly, the guidelines did not address potential effects on mental health.

Preclinical studies demonstrate that aspartame consumption is associated with neurobehavioral changes including anxiety-like behavior and learning and memory deficits ^24, 30–35^. A recent report by us ^35^ demonstrated that aspartame-induced anxiety-like behavior is heritable along the paternal line of descent. However, whether cognitive deficits associated with aspartame consumption such as learning, and memory deficits can be heritable was not known. Moreover, traditionally the risk of adverse health effects in future generations is appreciated in the context of deleterious environmental exposures of pregnant and nursing women. Environmental exposures of men are not considered significant in this context.

Here we use a mouse model and report that aspartame consumption via drinking water daily for up to 16 weeks at doses equivalent to only 7-15% of the FDA recommended maximum daily intake value for humans [roughly equivalent to human consumption of 2-4 small (8 oz) diet soda drinks per day] produces significant deficits in spatial learning and memory. Perhaps even more interestingly, the learning and memory deficits are transmitted by aspartame consuming male mice to their male and female offspring.

## Results

### A mouse model of aspartame exposure

Adult (approximately 8-week-old) C57BL/6 male mice were given free access to drinking water containing aspartame (0.03% or 0.015%), while another set of male mice received plain drinking water (control group; Fig. 1). Our previous work ^35^ showed that C57BL/6 mice showed neither a preference nor aversion to drinking water containing 0.03% or 0.015% aspartame ^35^. In addition, consumption of drinking water containing 0.015% or 0.03% aspartame daily for up to 18-weeks did not produce significant effects on bodyweight or serum metabolic markers ^35^.

**Figure 1:**
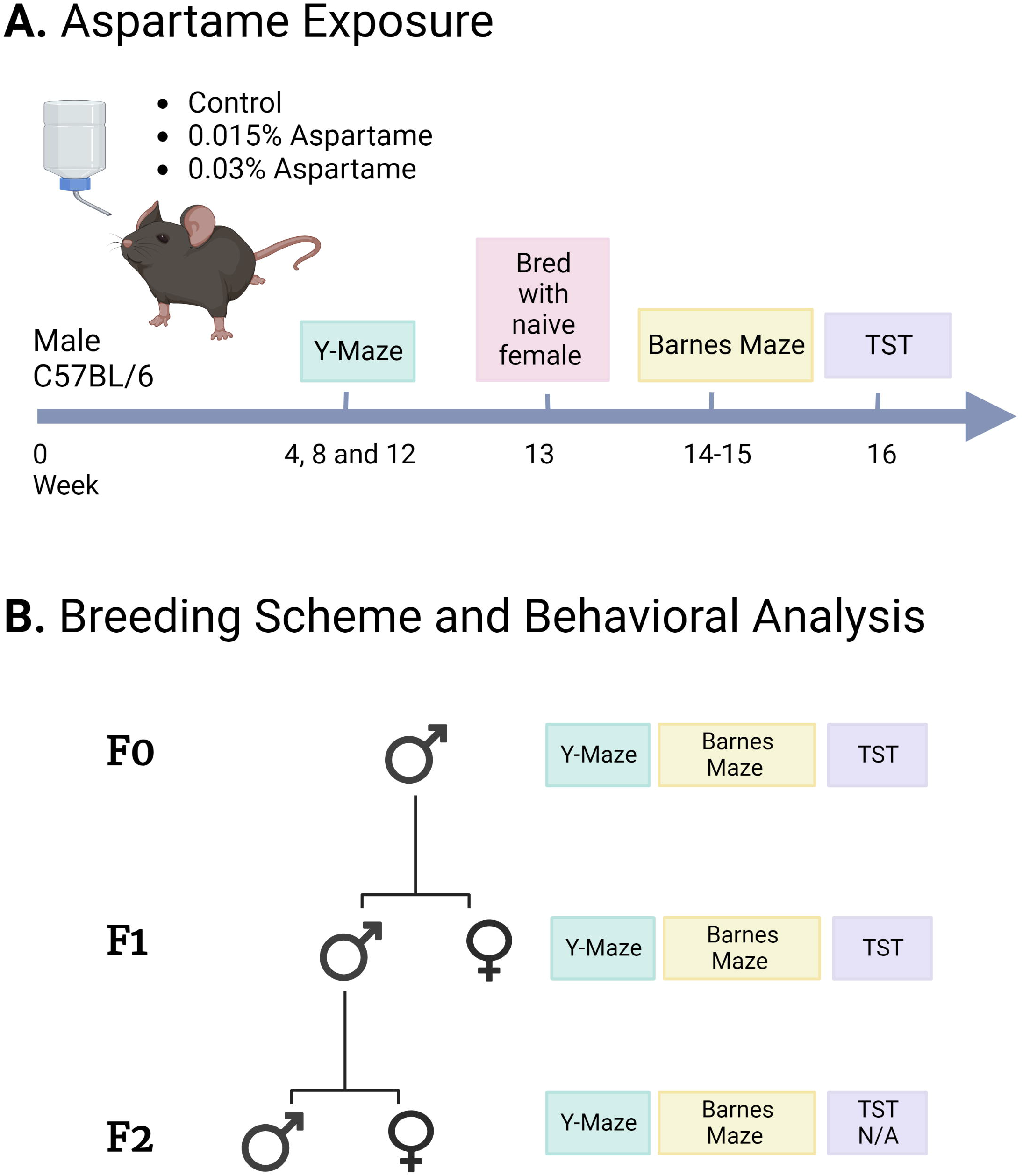
Schematic representation of the experimental design. (A) Male C57BL/6 mice were exposed to drinking water containing 0.015% or 0.03% aspartame for up to 16 weeks. Control groups received plain drinking water over the same duration. Y-maze assay was performed at 4, 8, and 12-weeks during the drinking water exposure. During the 13^th^ week, the mice were bred with females purchased from the vendor and maintained on plain drinking water to produce the F1 generation. Following the breeding, and during weeks 14-15 of the exposures, Barnes maze assay was performed. During the 16^th^ week, tail suspension test (TST) was performed. (B). Beginning at approximately 2-months of age, the F1 male and female mice from each drinking water lineage completed the Y-maze, Barnes maze and TST. Male mice from the 0.03% F1 lineage that did not undergo behavioral testing but were littermates of F1 males that did perform behavioral tests, were bred with females purchased from the vendor and maintained on plain drinking water to produce the F2 generation. Y-maze, and Barnes maze assays (but not TST) were performed in F2 male and female mice beginning at approximately 2 months of age. This Figure was created with BioRender.com.

The average body weight of the adult male mice in the present study was 26.0 g. Our previous work ^35^ showed that adult male C57BL/6 mice consumed approximately 7.0 ml of plain drinking water daily. Therefore, on average, male mice in the 0.03% and 0.015% aspartame groups consumed 86.4 mg/kg and 43.2 mg/kg aspartame per day, respectively ^35^. Based on allometric conversion utilizing pharmacokinetic and body surface area parameters ^36^, the daily aspartame dose delivered to male mice in the two aspartame groups was 7-15% of the FDA recommended human maximum daily intake value of 50 mg/kg ^25, 28^. It is estimated that the actual average daily aspartame consumption in humans is only 4.1 mg/kg ^37^, well below FDA’s maximum daily intake value. In the present study, mice consuming 0.015% aspartame received the mouse equivalent of the 4.1 mg/kg human dose ^36^.

During the 16 weeks of aspartame exposure, we performed tests of spatial working memory (Y-maze), spatial learning, memory, and reversal learning (Barnes maze) and learned helplessness (tail suspension test, TST; Fig. 1).

Since heritability of behavioral traits via the paternal line of descent has not been examined extensively, we examined male mice rather than female mice in the directly exposed generation, as well as their male and female descendants (Fig. 1).

### Spatial working memory

The Y-maze assay was performed every four weeks over the 12 weeks of aspartame containing or plain drinking water exposure (i.e., at weeks 4, 8 and 12; Fig. 1). The percent spontaneous alternations, a measure of spatial working memory, showed significant effect of drinking water treatment (Fig 2A; Repeated Measures Two-way ANOVA; F_(2,21)_ = 14.35, p<0. 0001; Table 1A) but not the duration of exposure or the interaction between drinking water treatment and duration of exposure (Table 1A). Thus, aspartame’s effects on working memory were present as early as 4 weeks of exposure and persisted over the entire 12-week duration. Although there were significant differences between the aspartame groups and control group in spontaneous alternations, there was no significant difference between the 0.015% and 0.03% aspartame groups at any of the 3 intervals (Table 1B).

**Figure 2:**
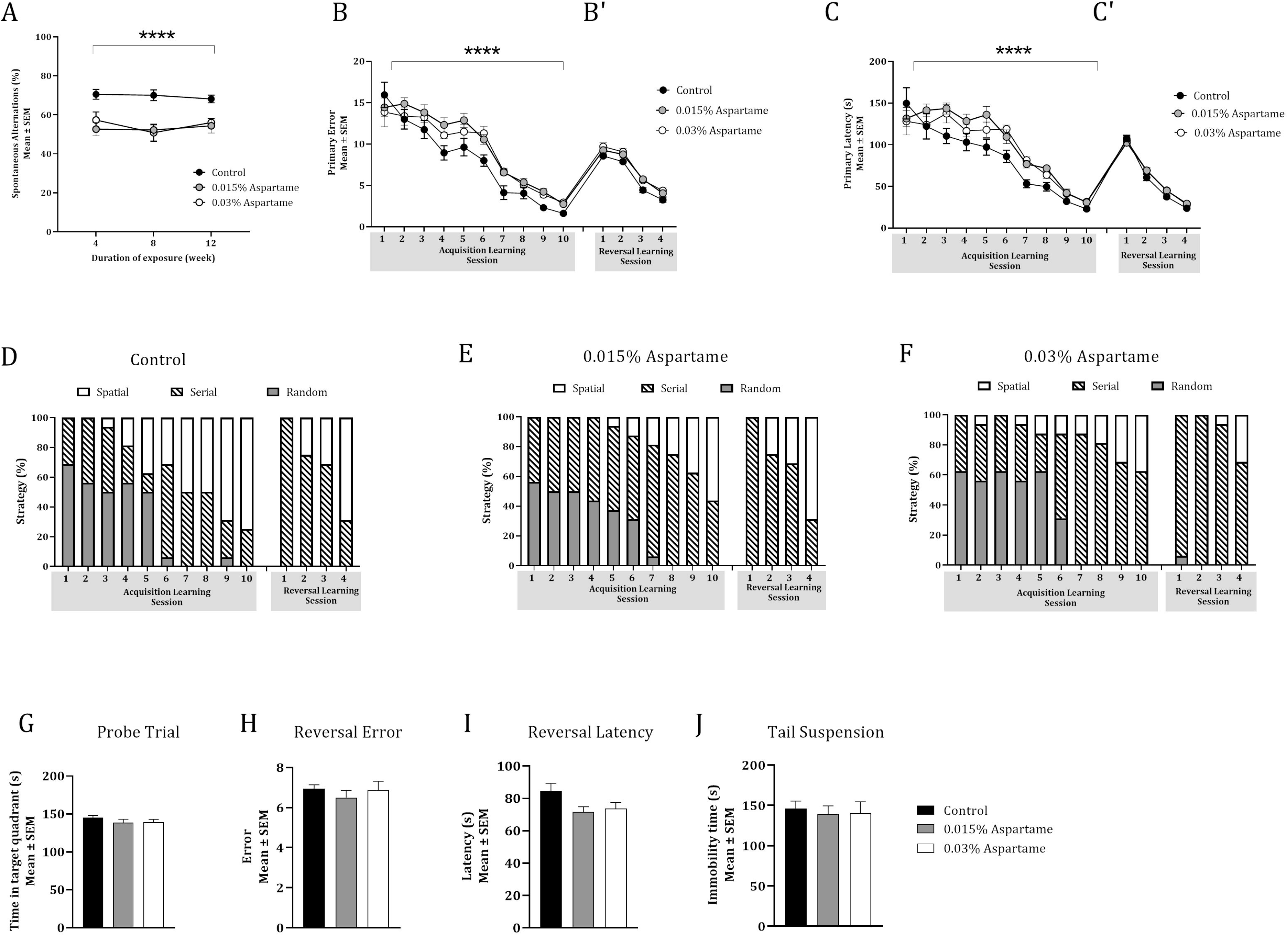
Spatial working memory, spatial learning and learned helplessness in male F0 mice. Spatial working memory was analyzed based on spontaneous alternations in a Y-maze (A) at 4, 8 and 12-weeks of the drinking water exposures. The drinking water exposure produced significant effects (***; Two-way Repeated Measures ANOVA, p<0.0001), suggesting poor performance by the two aspartame groups. Spatial learning was analyzed in the Barnes maze based on primary errors (B) and primary latency (C) during 10 consecutive daily acquisition of learning sessions and 4 daily consecutive reversal learning sessions (B’ and C’). Drinking water exposure produced significant effects on primary errors and primary latency during acquisition of learning (****; Two-way Repeated Measures ANOVA, p<0.0001), as did the sessions. Drinking water exposure did not produce significant effects on primary errors or primary latency (B’ an C’) but the sessions did, on both measures. The three types of search strategy employed during each session of acquisition of learning and reversal learning were analyzed for the control (D), the 0.015% aspartame (E) and 0.03% aspartame (F) groups. All three groups transitioned from random or serial strategies to predominantly spatial strategy (open bars) by the final sessions of acquisition (session #10) and reversal learning (session #4). Spatial memory retention/recall based on the time spent in the target quadrant during the probe trial (G) did not show significant effects of drinking water treatment. Similarly, the reversal effect based on primary errors (H) or primary latency (J) did not show significant effects of drinking water exposure. Drinking water exposure did not produce significant effects on the total time immobile during the tail suspension test (J).

**Table 1:** Statistical analyses of the data for male mice in the founder (F0) generation consuming drinking water containing 0.015% or 0.03% aspartame or plain drinking water (control). Table 1: Statistical analyses of the data from Y-maze (A-C), Barnes maze (D-L) and tail suspension test (M) for male mice in the founder (F0) generation that consumed drinking water containing aspartame (0.015% or 0.03%) or plain drinking water (control). The statistical test used for each analysis is indicated. Spatial working memory was analyzed using spontaneous alternations (A,B) and total number of arm entries (C) in the Y-maze. Spatial learning and memory [acquisition of learning (D, E, F), probe trial (G) and reversal learning (H-L)) were analyzed using the Barnes maze. Learned helplessness was analyzed using the tail suspension test (M). Statistical significance is indicated as follows: * = p<0.05; ** = p<0.01; *** = p<0.001; and **** = p<0.0001.

| Behavioral Parameter and Statistical Test |  | Variable | F<sub>(df)</sub> (ANOVA)<br>t (Bonferroni)<br>χ<sup>2</sup><sub>(df)</sub> (Chi-square) |  |
| --- | --- | --- | --- | --- |
| A | Spontaneous alternations in Y-maze<br>(Repeated Measures Two-way ANOVA) | Drinking water treatment | 14.35<sub>(2,21)</sub> **** |  |
|  |  | Duration of exposure | 0.78<sub>(1.43, 30.09)</sub> |  |
|  |  | Treatment x Duration | 0.77<sub>(4,42)</sub> |  |
| B | Spontaneous alternations in Y-maze<br>(Bonferroni Multiple Comparisons for each of 3 intervals) | 0.015% aspartame versus Control | Week 4 | 4.23** |
|  |  |  | Week 8 | 4.78*** |
|  |  |  | Week 12 | 3.27* |
|  |  | 0.03% aspartame versus Control | Week 4 | 2.73 |
|  |  |  | Week 8 | 3.72** |
|  |  |  | Week 12 | 4.30** |
|  |  | 0.015% versus 0.03% aspartame | Week 4 | 0.87 |
|  |  |  | Week 8 | 0.28 |
|  |  |  | Week 12 | 0.39 |
| C | Total number of arm entries in Y-maze<br>(Repeated Measures Two-way ANOVA) | Drinking water treatment | 0.12<sub>(2,21)</sub> |  |
|  |  | Duration of exposure | 0.78<sub>(1.58, 33.26)</sub> |  |
|  |  | Treatment x Duration | 0.18<sub>(4,42)</sub> |  |
| D | Barnes Maze, Acquisition of Learning: Primary errors<br>(2-way Repeated Measures ANOVA) | Session | 88.83<sub>(4.18,87.75)</sub> **** |  |
|  |  | Drinking water treatment | 14.07<sub>(2,21)</sub> *** |  |
|  |  | Session x | 1.11<sub>(18,189)</sub> |  |

**Table 1:** Statistical analyses of the data from Y-maze (A-C), Barnes maze (D-L) and tail suspension test (M) for male mice in the founder (F0) generation that consumed drinking water
|  |  | Treatment |  |
| --- | --- | --- | --- |
| E | Barnes Maze, Acquisition of Learning: Primary latency<br>(2-way Repeated Measures ANOVA) | Session | 61.64<sub>(4.64,97.40)</sub> **** |
|  |  | Drinking water treatment | 10.63<sub>(2,21)</sub> *** |
|  |  | Session x Treatment | 1.24<sub>(18,189)</sub> |
| F | Search Strategy during Acquisition of Learning<br>(Chi-square) | Drinking water treatment | 30.96<sub>(14)</sub> ** |
| G | Barnes Maze, Probe Trial: Time in target quadrant (1-way ANOVA) | Drinking water treatment | 0.86<sub>(2,21)</sub> |
| H | Barnes Maze, Reversal Effect: Primary errors (1-way ANOVA) | Drinking water treatment | 0.47<sub>(2,21)</sub> |
| I | Barnes Maze, Reversal Effect: Primary latency (1-way ANOVA) | Drinking water treatment | 0.76<sub>(2,21)</sub> |
| J | Barnes Maze, Reversal Learning over 4 days: Primary errors (2-way Repeated Measures ANOVA) | Session | 213.60<sub>(2.34,49.06)</sub> **** |
|  |  | Drinking water treatment | 2.86<sub>(2,21)</sub> |
|  |  | Session x Treatment | 0.46<sub>(6,63)</sub> |
| K | Barnes Maze, Reversal Learning over 4 days: Primary latency (2-way Repeated Measures ANOVA) | Session | 389.30<sub>(2.51,52.79)</sub> **** |
|  |  | Drinking water treatment | 2.79<sub>(2,21)</sub> |
|  |  | Session x Treatment | 1.23<sub>(6,63)</sub> |
| L | Search Strategy during Reversal Learning<br>(Chi-square) | Drinking water treatment | 27.87<sub>(14)</sub> **** |
| M | Total time immobile in Tail Suspension test<br>(One-way ANOVA) | Drinking water treatment | 0.10<sub>(2,21)</sub> |

The total number of arm entries, a proxy for general locomotion and maze exploration did not show significant effects of drinking water treatment, duration of exposure, or interaction between drinking water treatment and duration of exposure (Repeated Measures Two-way ANOVA; Table 1C). Therefore, the differences in percent spontaneous alternations among the drinking water groups were not due to significant differences in the ability of the mice to perform the Y-maze test.

### Spatial learning and memory

Next, we used the Barnes maze to evaluate spatial learning and memory ^38–41^. Male mice were tested beginning at 14 weeks of the drinking water exposure paradigm (Fig. 1). The test began with acquisition of learning phase with two acquisition trials during each daily session and continued over 10 consecutive sessions (days). During each acquisition trial, the time taken to locate the escape box (i.e., primary latency) and the number of errors committed prior to locating the escape box (primary errors) were recorded ^42, 43^. Primary latency and primary errors are sensitive measures of group differences compared to total latency (length of time taken to locate the escape box and then to enter it) and total number of errors (number of errors committed before and after locating the escape box) ^42^. The data from the 2 trials were averaged and represented as data for that session.

Primary error data showed significant effects of session (Fig. 2B; Repeated Measures Two-way ANOVA; F_(4.18, 87.75)_ = 88.83, p<0.0001; Table 1D) and drinking water treatment (F_(2,21)_ = 14.07, p<0.001; Table 1D) but no significant interaction between the two (Table 1D). Primary latency data also showed significant effects of session (Fig. 2C; Repeated Measures Two-way ANOVA; F_(4.64, 97.40)_ = 61.64, p<0.0001; Table 1E) and the drinking water treatment (Repeated Measures Two-way ANOVA; F_(2,21)_ = 10.63, p<0.001; Table 1E), but no significant interaction between the two (Table 1E). Thus, although the performance of the mice in all three drinking water groups improved with each successive session (i.e., primary errors and primary latency decreased significantly with each successive session), the performance of the mice exposed to aspartame (0.03% or 0.015%) improved slowly compared to that of the mice exposed to plain drinking water (Fig. 2B, C). However, the performance of the mice in the two aspartame groups (0.015% and 0.03%) was not significantly different from each other. Bonferroni pair-wise comparisons showed that the primary errors in mice in the 0.015% and 0.03% aspartame groups were significantly greater than the mice in the control group during sessions 6 and 9 whereas the 0.015% aspartame group had significantly more primary errors during session 4 as well (Supplementary Table 2C). Compared to the control group, the primary latency was significantly greater in the 0.015% aspartame group during sessions 3, 5, 7 and 9, whereas it was significantly greater in the 0.03% aspartame group during sessions 6 and 7 (Supplementary Table 2C). There was no significant difference between the two aspartame groups in primary errors or latency during any of the 10 sessions.

Mice use spatial, serial, or random search strategies to find the escape box. The spatial strategy involves moving directly to the escape box or to the 2 holes on either side of it. The serial strategy involves visiting at least 3 holes adjacent to the escape box in a serial manner before visiting the escape box and crossing the center of the arena (i.e., the starting point) only one time. The random strategy involves unorganized search patterns including crossing the center more than once. Since the position of the escape box remained invariant over successive trials and sessions, and since we used distant spatial cues positioned on the walls of the room (rather than local or intra-maze cues) the adoption of the spatial search strategy indicates utilization of spatial cues to learn the location of the escape box ^42, 43^. In other words, the time taken (i.e., number of sessions needed) to adopt the spatial search strategy is a reliable measure of spatial learning and memory.

Mice in all 3 groups used random or serial strategies and not the spatial strategy during session #1 of acquisition of learning, but gradually transitioned to the spatial strategy (Fig. 2D-F). The time taken to successfully transition to the spatial strategy was significantly different among the three groups (Fig. 2G; χ^2^ =30.96, p<0.01; Table 1F). Thus, although there was a progressive shift toward the spatial search strategy in all 3 drinking water groups, the aspartame groups lagged the plain drinking water group in making this transition during the acquisition of learning phase, suggesting a deficit in spatial learning.

Following the 10 sessions of acquisition of learning, on the 11^th^ day of the Barnes maze test, an acquisition probe trial was performed to assess retention and retrieval of spatial memory. During this trial, the escape box was removed from the arena. In this paradigm, mice with intact retention and retrieval of spatial memory revert to their previously learned behavior and selectively focus their search on the previous location of the escape box. In other words, they spend longer proportion of their search in the quadrant representing the former location of the escape box (target quadrant) compared to the other quadrants of the arena ^42, 43^.

The probe trial data showed that drinking water treatment did not produce significant effects on the time spent in the target quadrant indicating that retention and retrieval of spatial memory was not affected by the aspartame treatment (Fig. 2G; Table 1G, One-way ANOVA).

The final phase of the Barnes maze assay evaluated reversal learning, which is a measure of cognitive flexibility. It is evaluated based on the ability to learn the location of the escape box under a new set of rules ^43–49^. Here, the rules are changed by moving the escape box from its original location to a new, diametrically opposite location. All other parameters remain the same as those in the previous 10 learning sessions. The primary latency to locate the escape box in its new location and the primary errors committed in the process are recorded over 4 consecutive sessions, two trials per session.

Mice in all 3 drinking water groups committed more primary errors (Fig. 2B’) and had greater primary latency (Fig. 2C’) during the first reversal learning session (#1) compared to the latency and errors during the final acquisition of learning session (#10), demonstrating the reversal effect. However, neither the primary errors nor primary latency contributing to the reversal effect was influenced significantly by the drinking water treatment (Fig. 2 H, I; One-way ANOVA, Table 1 H, I). Thus, the aspartame exposure did not produce significant effects on reversal learning.

When the latency to find the escape box in its new position and the number of primary errors committed in the process were analyzed across the full complement of the 4 reversal learning sessions, both measures showed a significant effect of session (Fig. 2 B’ and C’; Repeated Measures Two-way ANOVA; primary errors, F_(2.34,49.06)_ = 213.60, p<0.0001; primary latency, F_(2.51,52.79)_ = 389.30, p<0.0001; Table 1 J, K) but no significant effects of drinking water treatment or treatment by session interaction (Table 1 J, K). Thus, the mice in all three drinking water groups re-learned the new location of the escape box under the reversed paradigm over the 4 reversal learning sessions, and the aspartame exposure did not produce significant effects on the re-learning.

Bonferroni pair-wise comparisons showed that the mice in the 0.015% and 0.03% aspartame groups committed significantly greater number of primary errors than the mice in the control group only during session 3 and there were no significant differences between the aspartame and control groups in primary latency during any of the 4 sessions (Supplementary Table 2E, F). There were no significant differences between the two aspartame groups in primary errors or latency during any of the 4 sessions (Supplementary table 2E, F).

Analysis of the search strategy during reversal learning showed that the mice in all three groups transitioned from serial or random strategies to spatial strategy over the 4 days of the reversal learning phase (Fig. 2 D-F). However, the time taken to successfully transition to the spatial strategy was significantly different among the three groups (χ^2^_(14)_ = 27.87, p<0.0001, Table 1L), suggesting a spatial learning deficit in the aspartame groups. Whether differences in search strategy are as robust a measure of spatial learning as are differences in primary errors or primary latency or if they reflect differences in the underlying brain circuitry reported in human learning disabilities remains to be examined ^42, 43, 50, 51^.

### Learned helplessness

We used the tail suspension test (TST) to evaluate learned helplessness in aspartame exposed and plain drinking water exposed male mice. TST has been used successfully in rodents to test the efficacy of human anti-depressant drugs, suggesting its utility as a test for depression-like behavior in rodents ^52–54^.

There was no significant effect of drinking water treatment on total time immobile (Fig 2J, One-way ANOVA; Table 1M). Thus, the aspartame exposure did not produce significant effects on learned helplessness behavior, and by extrapolation, on depression-related behavior.

### Heritability of the behavioral effects: F1 generation

We bred male mice that were exposed to 0.015% or 0.03% aspartame-containing drinking water (F0 generation) for 12-weeks with female mice purchased from the vendor maintained on plain drinking water to produce the F1 generation (Fig. 1). Male mice in the plain drinking water control group were bred with female mice purchased from the vendor and maintained on plain drinking water to generate control F1 generation (Fig. 1) The F0 male mice from the two aspartame and the control groups had completed Y-maze assay but not the Barnes maze or TST prior to breeding with the females (Fig. 1). The latter tests were performed after the breeding.

We analyzed litter size and developmental milestones in all three drinking water groups from the F1 generation (Supplementary Table 1). The drinking water treatment did not produce significant effects on any of the measurements, suggesting that aspartame consumption did not affect pregnancy outcomes or offspring growth and development.

Heritability via the maternal lineage was not the focus of this study. Historically, the effects of environmental factors on subsequent generations are considered following exposure of females, especially pregnant and nursing mothers ^55–61^. The impact of paternal environmental exposures on heritable traits has not attracted as much attention ^38^. Therefore, the present study focused on heritability along the paternal lineage.

### Spatial working memory in the F1 generation

The percent spontaneous alternations in the Y-maze showed a significant effect of paternal lineage (Fig 3A; Mixed model ANOVA; F_(2,33)_ = 27.72, p<0.0001; Table 2A), but the effect of sex or the interaction between paternal lineage and sex were not significant (Fig 3A; Table 2A). Post hoc contrast analysis showed significant deficits in spontaneous alternation in the 0.015% and 0.03% aspartame lineages compared to the control lineage (0.015% versus control, t_(2,33)_ = 6.24; p<0.0001; 0.03% versus control, t_(2,33)_ = 6.64, p<0.0001; Supplementary Table 2). The difference between the two aspartame lineages was not significant (Supplementary Table 2). These data demonstrate significant spatial working memory deficit in F1 male and female mice derived from both the aspartame lineages compared to the F1 mice from the control lineage.

**Figure 3:**
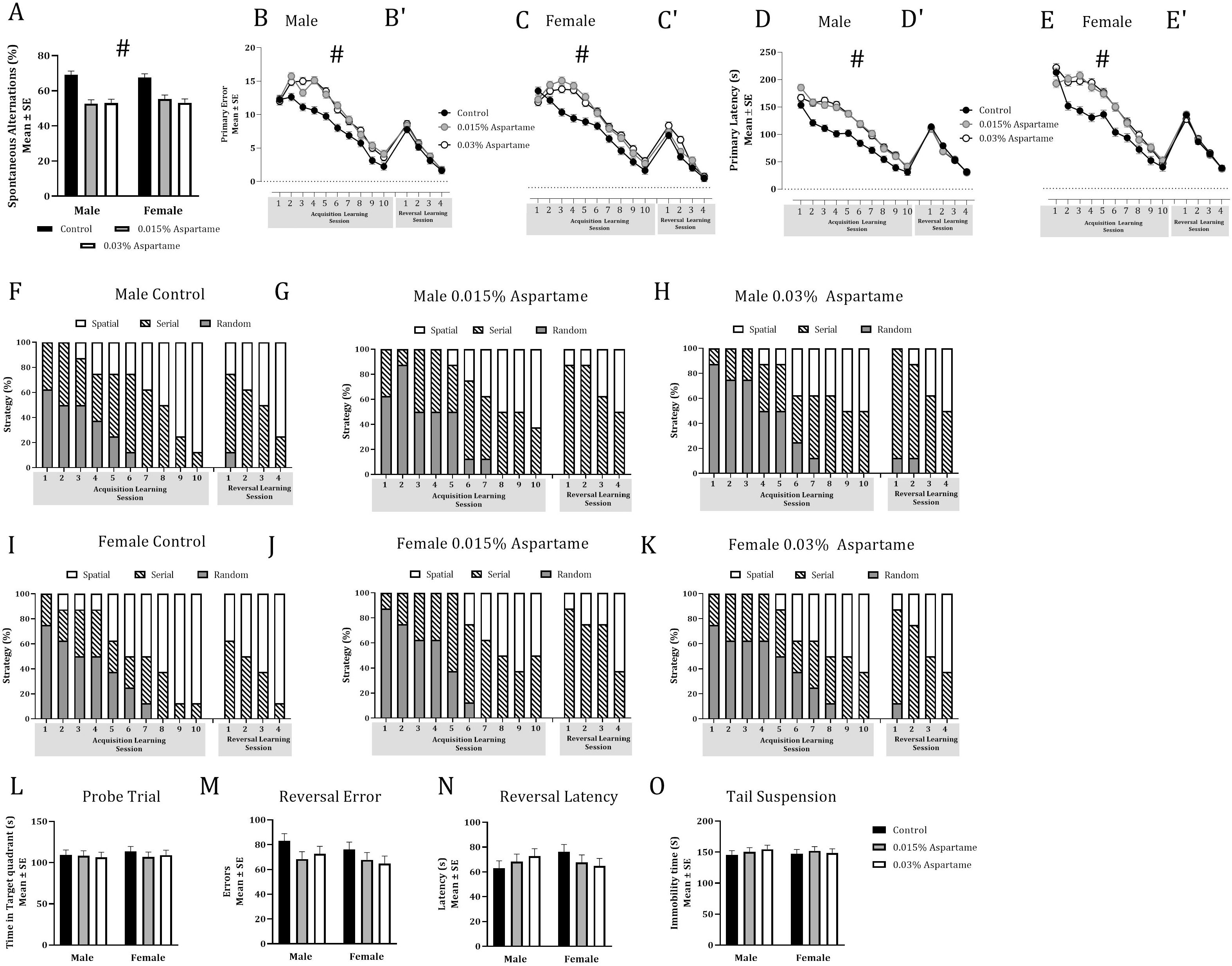
Spatial working memory, spatial learning and learned helplessness in male and female F1 mice. Mice from the 0.015% and 0.03% aspartame lineages showed significant deficits in spontaneous alternation in the Y-maze compared to the mice from the control group (A; #; Mixed model ANOVA, p<0.0001). Spatial learning was analyzed in the Barnes maze based on primary errors (Male: B, B’; Female: C, C’) and primary latency (Male: D, D’; Female: E, E’) during 10 daily acquisition of learning sessions and 4 daily reversal learning sessions. Sessions and lineage each produced significant effects on primary errors (B, C) and primary latency (D, E) during acquisition of learning phase (# Mixed model ANOVA; p<0.0001). Males (F-H) and females (I-K) in all lineages transitioned from random or serial strategies to predominantly spatial strategy during acquisition of learning and reversal learning. However, male (F-H) and female (I-K) mice in the aspartame lineages lagged the mice in the control lineage in the transition during the acquisition of learning phase (Chi-square; p<0.01). Time spent in the target quadrant during the probe trial (L) did not show significant effects of drinking water treatment. Similarly, the reversal effect based on a comparison of the primary error (M) or primary latency (N) between acquisition of learning session #10 and reversal learning session #1 did not show significant effects of drinking water treatment. Drinking water exposure did not produce significant effects on the total time immobile in the tail suspension test (O).

**Table 2:** Statistical analyses of the data for male and female mice in the F1 and F2 generations derived from the paternal aspartame or plain drinking water (control) lineages. Table 2: Statistical analyses of the data for male and female mice in the F1 and F2 generations derived from the paternal aspartame or plain drinking water (control) lineages. The F1 mice were derived from two paternal aspartame lineages (0.015% and 0.03%) whereas the F2 mice were from a single aspartame lineage (0.03%). A mixed model ANOVA with litter as random factor was used. Spatial working memory was analyzed using spontaneous alternations (A) and total number of arm entries (B) in the Y-maze. Spatial learning and memory [acquisition of learning (C, D), probe trial (E) and reversal learning (F-K)] were analyzed using the Barnes maze. Learned helplessness was analyzed using the tail suspension test (L) in the F1 generation only. Statistical significance is indicated as follows: ** = p<0.01 and **** = p<0.0001.

| Behavioral Parameter and Statistical Test |  | Variable | F<sub>(df)</sub> Mixed Model ANOVA<br>t<sub>(df)</sub> Post hoc test<br>χ<sup>2</sup><sub>(df)</sub> Chi Square |  |
| --- | --- | --- | --- | --- |
|  |  |  | F1 | F2 |
| A | Spontaneous alternations in Y-maze (ANOVA) | Lineage | 27.72<sub>(2, 33)</sub> **** | 0.1<sub>(1,22)</sub> |
|  |  | Sex | 0.06<sub>(1,33)</sub> | 2.91<sub>(1,22)</sub> |
|  |  | Lineage x sex | 0.52<sub>(2, 33)</sub> | 0.98<sub>(2,22)</sub> |
| B | Total number of arm entries in Y-maze (Two-way ANOVA) | Lineage | 1.21<sub>(2,33)</sub> | 0.25<sub>(1,22)</sub> |
|  |  | Sex | 0.18<sub>(2,33)</sub> | 0.14<sub>(1,22)</sub> |
|  |  | Lineage x sex | 0.29<sub>(2,33)</sub> | 0.06<sub>(2,22)</sub> |
| C | Barnes Maze, Acquisition of Learning: Primary errors (ANOVA) | Session | 287.35<sub>(2,171)</sub> **** | 433.31<sub>(9,114)</sub> **** |
|  |  | Lineage | 103.99<sub>(2,171)</sub> **** | 0.35<sub>(1,114)</sub> |
|  |  | Sex | 0.12<sub>(1,171)</sub> | 0.14<sub>(1,114)</sub> |
|  |  | Session x lineage | 4.38<sub>(18,171)</sub> **** | 0.46<sub>(9,114)</sub> |
|  |  | Session x sex | 1.04<sub>(9,141)</sub> | 0.71<sub>(9,114)</sub> |
|  |  | Lineage x sex | 0.89<sub>(1,171)</sub> * | 0.65<sub>(1,114)</sub> |
|  |  | Session x lineage x sex | 0.86<sub>(18,171)</sub> | 0.95<sub>(9,114)</sub> |
| D | Barnes Maze, Acquisition of Learning: Primary latency (ANOVA) | Session | 303.5<sub>(9,171)</sub> **** | 354.07<sub>(9,114)</sub> **** |
|  |  | Lineage | 142.65<sub>(2,171)</sub> **** | 0.01<sub>(1,114)</sub> |
|  |  | Sex | 0.01<sub>(2,171)</sub> | 4.61<sub>(1,114)</sub> * |
|  |  | Session x lineage | 3.78<sub>(18,171)</sub> **** | 0.70<sub>(9,114)</sub> |
|  |  | Session x sex | 0.12<sub>(9,171)</sub> | 1.20<sub>(9,114)</sub> |
|  |  | Lineage x sex | 1.28<sub>(2,171)</sub> | 0.16<sub>(1,114)</sub> |
|  |  | Session x lineage x sex | 1.03<sub>(18,171)</sub> | 0.23<sub>(9,114)</sub> |
| E | Transition to Spatial Search Strategy during Acquisition of Learning (Chi-square) | Male | 52.74<sub>(14)</sub>**** | 36.36<sub>(7)</sub>**** |
|  |  | Female | 61.75<sub>(14)</sub>**** | 10.17<sub>(7)</sub> |
| F | Barnes Maze, Probe Trial: Time in target quadrant (ANOVA) | Lineage | 0.23<sub>(2,9)</sub> | 1.48<sub>(1,6)</sub> |
|  |  | Sex | 0.13<sub>(1,9)</sub> | 0.01<sub>(1,6)</sub> |
|  |  | Lineage x sex | 0.13<sub>(2,9)</sub> | 0.14<sub>(1,6)</sub> |
| G | Barnes Maze, Reversal Effect: Primary errors (ANOVA) | Lineage | 1.86<sub>(2,9)</sub> | 0.2<sub>(1,6)</sub> |
|  |  | Sex | 1.48<sub>(1,9)</sub> | 0.0<sub>(1,6)</sub> |
|  |  | Lineage x sex | 0.29<sub>(2,9)</sub> | 2.35<sub>(1,6)</sub> |
| H | Barnes Maze, Reversal Effect: Primary latency (ANOVA) | Lineage | 1.86<sub>(1,9)</sub> | 2.89<sub>(1,6)</sub> |
|  |  | Sex | 1.46<sub>(1,9)</sub> | 0.02<sub>(1,6)</sub> |
|  |  | Lineage x sex | 0.29<sub>(1,9)</sub> | 1.50<sub>(1,6)</sub> |
| I | Barnes Maze, Reversal Learning - 4 sessions: Primary errors (ANOVA) | Session | 266.95<sub>(3,63)</sub>**** | 131.49<sub>(2,30)</sub>**** |
|  |  | Lineage | 1.86<sub>(2,63)</sub> | 0.05<sub>(1,30)</sub> |
|  |  | Sex | 0.19<sub>(1,63)</sub> | 1.30<sub>(1,30)</sub> |
|  |  | Session x lineage | 1.51<sub>(6,63)</sub> | 0.24<sub>(2,30)</sub> |
|  |  | Session x sex | 0.2<sub>(3,63)</sub> | 0.70<sub>(1,30)</sub> |
|  |  | Lineage x sex | 1.01<sub>(2,63)</sub> | 1.30<sub>(1,30)</sub> |
|  |  | Session x lineage x sex | 0.88<sub>(6,63)</sub> | 0.70<sub>(2,30)</sub> |
| J | Barnes Maze, Reversal Learning - 4 sessions: Primary latency (ANOVA) | Session | 479.01<sub>(3,63)</sub>**** | 605.42<sub>(2,30)</sub>**** |
|  |  | Lineage | 0.23<sub>(2,63)</sub> | 0.20<sub>(1,30)</sub> |
|  |  | Sex | 4.38<sub>(1,63)</sub>* | 0.71<sub>(1,30)</sub> |
|  |  | Session x lineage | 0.23<sub>(6,63)</sub> | 1.60<sub>(2,30)</sub> |
|  |  | Session x sex | 0.48<sub>(3,63)</sub> | 7.22<sub>(1,30)</sub>*** |
|  |  | Lineage x sex | 0.61<sub>(2,63)</sub> | 0.40<sub>(1,30)</sub> |
|  |  | Session x lineage x sex | 0.87<sub>(6,63)</sub> | 2.02<sub>(2,30)</sub> |
| K | Transition to Spatial Search Strategy during Reversal Learning (Chi-square) | Male | 3.94<sub>(4)</sub> | 2.52 <sub>(2)</sub> |
|  |  | Female | 8.01<sub>(4)</sub> | 3.94 <sub>(2)</sub> |
| L | Total time immobile in Tail Suspension Test (ANOVA) | Lineage | 0.27 <sub>(2,33)</sub> | N/A |
|  |  | Sex | 0.03 <sub>(1,33)</sub> |  |
|  |  | Lineage x sex | 0.28 <sub>(2,33)</sub> |  |

Analysis of total number of arm entries did not show significant effects of paternal lineage, sex, or the interaction between paternal lineage and sex (Table 2B; Supplementary Table 2). Therefore, the deficit in spatial working memory in the paternal aspartame lineages was not attributable to any change in the ability of the mice to navigate the Y-maze.

### Spatial learning and memory in the F1 generation

Analysis of the acquisition of learning data in the Barnes maze showed that the number of primary errors committed (Fig. 3B, C) had significant effects of session (F_(9,171)_ = 287.35, p<0.0001; Table 2C) and paternal lineage (F_(2,171)_ = 103.99, p<0.0001; Table 2C), but the effect of sex was not significant (Table 2 C). There was a significant interaction between session and paternal lineage (F_(18,171)_ = 4.38, p<0.0001; Table 2C) and lineage and sex (F_(1,171)_ = 0.89, p<0.05; Table 2C). However, none of the other interactions between session, sex and treatment were significant (Table 2 C). A post hoc contrast analysis revealed that the differences between 0.015% aspartame lineage and control lineage and 0.03% aspartame lineage and control lineage were significant during each of the 10 acquisition sessions except on session #1 (Supplementary Table 2C). The differences between the two aspartame lineages were not significant for any of the 10 sessions (Supplementary Table 2C). Thus, both aspartame lineages showed significant acquisition of learning deficits compared to the control lineage and the deficits were present throughout sessions 2-10.

The data on the primary latency to find the escape box (Fig. 3D, E); showed differences comparable to those observed in primary errors. The primary latency showed a significant effect of session (F_(9,171)_ = 303.5, p<0.0001; Table 2D) and paternal lineage (F_(2,171)_ = 142.65, p<0.0001; Table 2D), but not sex (Table 2D). The interaction between session and paternal lineage was significant (F_(18,171)_ = 3.78, p<0.0001; Table 2D). However, none of the other interactions between session, sex, and paternal lineage was significant (Table 2D). The differences between the 0.015% aspartame lineage and control lineage and between the 0.03% aspartame lineage and control lineage were significant during sessions 2-9 (Supplementary Table 2D). The differences between the two aspartame lineages were not significant for any of the 10 sessions (Supplementary Table 2D)). Thus, both the aspartame lineages showed significant acquisition of learning deficits compared to the control lineage and the deficits were present throughout sessions 2-9.

In summary, both male and female F1 mice derived from the 0.015% and 0.03% aspartame paternal lineages showed deficits in spatial learning compared to their counterparts in the plain drinking water paternal lineage. Therefore, the spatial learning deficits observed in the F0 male mice exposed directly to aspartame were transmitted to their male and female descendants in the F1 generation.

Male and female mice in both the lineages began the acquisition of learning sessions with random or serial search strategies (Fig. 3F-K) and transitioned to spatial search strategy with each successive acquisition of learning session. However, the F1 mice from the aspartame lineages lagged those from the plain water lineage in making this transition (Fig. 3 F-K). The difference between the lineages was significant (Chi-square, Male: χ2_(14)_ = 52.74, p<0.0001; Female:χ2_(14)_ = 61.75, p<0.0001; Table 2E). Thus, the deficits in spatial learning observed in the F0 aspartame male founders exposed to aspartame were inherited by their descendants in the F1 generation.

Analysis of data on the time spent in the target quadrant during the probe trial (Fig. 3L) did not show significant effects of paternal lineage, sex, or interactions between paternal lineage and sex (Table 2F). Thus, aspartame lineage did not produce significant effects on spatial memory retention or retrieval in male or female mice in the F1 generation.

The reversal effect was calculated as the difference in the number of primary errors or the primary latency between acquisition of learning session #10 and reversal learning session #1. Mice in all 3 lineages had greater primary latency and committed greater number of primary errors during the first reversal learning session (Reversal session #1) compared to the latency and errors during the final acquisition of learning session (Acquisition session #10; Fig. 3B’-E’), demonstrating the reversal effect. However, the difference in primary latency and primary errors contributing to the reversal effect was not influenced by lineage (Fig. 3 M, N; Mixed model ANOVA, Table 2 G-H). Thus, the mice in all three lineages showed the reversal effect.

Analysis of the primary error data during the 4 reversal learning sessions showed a significant effect of session (F_(3,63)_ = 266.95, p<0.0001; Table 2I). Neither the lineage nor sex showed significant effects (Table 2I). A significant difference between the 0.03% aspartame and control lineage was found during session #2 (Supplementary Table 2E). The differences between 0.015% aspartame and control lineages or between the two aspartame lineages were not significant (Supplementary Table 2E). The primary latency data showed significant effects of session (F_(3,63)_ = 479.01, p<0.0001; Table 2J) and sex (F_(1,63)_ = 4.38, p<0.05; Table 2J). The effects of lineage were not significant (Table 2J). The interactions between session, lineage and sex were not significant (Table 2J). Post hoc analysis did not reveal significant differences between the lineages during any of the 4 sessions (Supplementary Table 2F). Thus, the mice in both the paternal lineages showed the reversal effect, but their paternal lineage did not influence reversal learning, a proxy for cognitive flexibility.

Analysis of the search strategy data showed that male and female mice from both the lineages began the reversal learning sessions with random and serial strategies and transitioned to the spatial strategy (Fig. 3 F-K). However, lineage did not produce significant effects on this transition (Chi-square, Male: χ2_(4)_ = 3.942, p>0.05; Female: χ2_(4)_ = 8.010, p>0.05; Table 2K).

Collectively, these data demonstrate that whereas mice derived from the aspartame paternal lineage showed a deficit in acquisition of learning, the aspartame lineage did not produce reversal learning deficits.

### Learned helplessness in the F1 generation

The TST data from the F1 generation showed that the total time immobile did not show significant effects of paternal lineage, sex, or the interaction between paternal lineage (Fig 3 O; Table 2L). Thus, the mice in the F1 generation from the 0.03% aspartame paternal lineage did not show significant changes in learned helplessness behavior, a proxy for depression-like behavior.

### Transgenerational heritability of the behavioral effects: F2 generation

The F1 male mice from 0.03% aspartame lineage and plain drinking water lineages were bred with females purchased from the vendor and maintained on plain drinking water to produce the F2 generation. Since the behavioral effects in the F1 mice derived from the 0.015% and 0.03% aspartame groups were comparable, we produced the F2 generation from mice in the 0.03% aspartame lineage only. A lack of behavioral changes in the F2 generation produced from the 0.015% aspartame F1 male lineage would have necessitated testing the F2 mice from the 0.03% F1 lineage, before ruling out transgenerational transmission. Therefore, analyzing F2 mice from the 0.03% F1 lineage first was the conservative approach.

The behavioral effects in the F1 generation (above) likely reflect the consequences of direct exposure of the founder (F0) spermatozoa to aspartame and epigenetic inheritance of the effects of aspartame ^3, 38, 61^. On the contrary, any behavioral changes in the F2 generation would represent transgenerational transmission ^3, 38, 61^ because neither the F2 generation nor the F1 spermatozoa that produced the F2 generation were directly exposed to aspartame at any time.

We analyzed litter size and developmental milestones in all three drinking water groups in the F2 generation (Supplementary Table 1). There was no significant effect of lineage on any of the measurements, suggesting that the aspartame lineage did not affect pregnancy outcomes or development of the offspring.

We did not find significant differences between the aspartame and plain water lineages in the F2 generation in spatial working memory (Y-maze), spatial learning, spatial memory retention, retrieval, or reversal learning (Barnes maze). The data are shown in Figure 4 and Table 2. Interestingly the F2 male but not female aspartame lineage lagged the F2 plain water lineage in transition to spatial search strategy during acquisition of learning phase (Fig. 4 F-I, Chi-square, male: χ^2^ = 36.36, p<0.001; Table 2E). However, paternal lineage did not produce significant effects on this transition during the reversal learning phase (Table 2 K).

**Figure 4:**
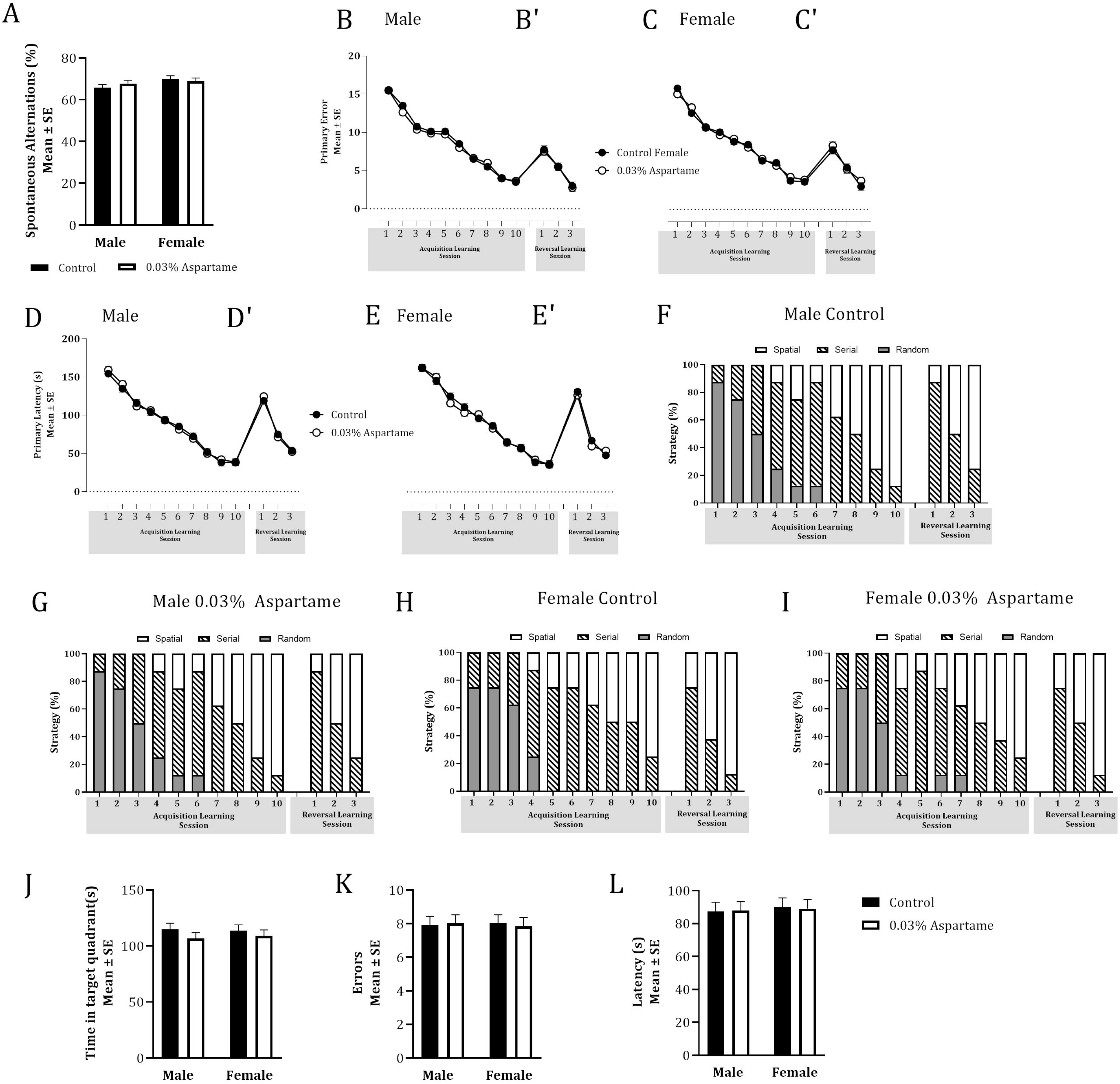
Spatial working memory and spatial learning in male and female mice in the F2 generation. Paternal lineage did not produce significant effects on spontaneous alternations in the Y-maze (A). During acquisition of learning in the Barnes maze, neither the primary errors (Male: B; Female: C)) not primary latency (Male: D; Female: E) showed significant effects of lineage, but the effects of sessions were significant (Mixed model ANOVA, p<0.0001). The reversal learning sessions (Male: B’, D’; Female: C’, E’) did not show significant effects of lineage but showed significant effects of session. Male (F,G) and female (H, I) mice from both the lineages transitioned from random or serial strategies to predominantly spatial strategy during acquisition of learning and reversal learning. Male mice in the aspartame lineage (G) lagged the male mice in the control lineage (F) in the transition during the acquisition of learning phase (Chi-square; p<0.01). Time spent in the target quadrant during the probe trial (J) did not show significant effects of lineage. Similarly, the reversal effect based on a comparison of the primary error (K) or primary latency (L) between acquisition of learning session #10 and reversal learning session #1 did not show significant effects of drinking water treatment.

There were no significant effects of 0.015% or 0.03% aspartame on learned helplessness in the F0 or F1 generations. Therefore, there was no empirical basis to expect changes in this behavior in the F2 generation. Therefore, we did not perform TST in the F2 generation.

## Discussion

We show that daily consumption of aspartame in drinking water by male mice at doses equivalent to 7-15% of the FDA recommended human maximum daily intake value [equivalent of the aspartame content of 2-4 cans of (small) 8 oz cans of diet soda] for up to 16 weeks produces significant deficits in spatial working memory and spatial learning. The spatial working memory deficit is apparent by 4 weeks of aspartame consumption, and persists until 12 weeks at least, the longest duration examined. Deficits in spatial learning were observed following 13 weeks of aspartame exposure. There were no deficits in spatial memory retention, recall or reversal learning. Aspartame consumption did not produce significant effects on learned helplessness, a proxy for depression-like behavior, even after 16 weeks. The spatial working memory and learning deficits were transmitted from the male mice to their male and female descendants in the F1 generation, demonstrating heritability of the traits but transgenerational transmission to the F2 generation was not observed.

Previous reports in preclinical models produced conflicting findings on the effects of aspartame consumption on learning and memory. Some reported deficits ^24, 30–34^, whereas others reported no effect ^62^ or even a “beneficial” effect ^63^. Clinical research produced similarly mixed results ^23, 64–69^. The lack of consensus may be due to differences in the experimental design including variability in the dose (<1 to 2000 mg/kg), relatively short duration of exposure (hours to 90 days) and variable routes of aspartame administration (oral, intraperitoneal, or subcutaneous). The inconsistent findings may have added to the controversy about aspartame’s CNS effects and may have contributed to prevailing notions about aspartame’s safety.

The present study used a dose, route and duration of aspartame exposure that represent long-term human aspartame use at only 7-15% of the FDA recommended maximum daily dose. Our findings not only demonstrate significant learning and memory deficits in male mice consuming aspartame, but also demonstrate a hitherto unrecognized consequence of aspartame consumption namely, heritability of the learning and memory deficits by male and female offspring in the F1 generation. A key public health implication of our findings is that the population at risk of aspartame’s adverse effects on learning and memory may be larger than current estimates, which consider the directly exposed individuals only. Our findings underscore the need for considering heritable effects as part of the safety evaluation of artificial sweeteners by regulatory agencies.

Historically, the consequences of parental deleterious exposures for their descendants were examined and understood in the context of exposure of the mothers, especially pregnant and nursing mothers ^55–61^. Our data show that exposure of the fathers is just as relevant when considering the mental health consequences for future generations.

The mechanisms underlying heritability of aspartame’s effects are not yet clear. Based on previous research on endocrine disrupting agents, nicotine, cocaine, alcohol, and cannabis^1, 3, 9, 12, 61, 70–76^ epigenetic changes in the spermatozoa emerge as candidate mechanisms. The alternative is genetic mechanisms such as mutations in spermatozoa, which would be expected to modify traits for generations. The effects of aspartame lasted only one generation, which would be consistent with epigenetic changes in the spermatozoa, which are transient and reversible ^77–82^.

The transience of environment-induced traits inherited via epigenetic changes in germ cells may fulfil the need for temporary adaptations to short-lived environmental insults and could have evolutionary significance. A permanent change mediated by mutations might not be beneficial and may be counterproductive to long-term adaptation if the environmental insult is short-lived.

We recently demonstrated that anxiety-like behavior produced by aspartame consumption shows transgenerational transmission along the paternal lineage because the anxiety persisted in the F2 generation ^83^. However, the magnitude of the anxiety attenuated during descent from F0 to F1 to F2 generations ^83^ indicating that even when transgenerational transmission of aspartame-induced traits is observed the magnitude of the effect is diluted from one generation to the next – as long as the aspartame exposure occurred only in the founder F0 generation. Similarly, behavioral traits produced by nicotine exposure of the founder generation show attenuation during descent from one generation to the next^3, 38, 61, 70, 84^.

Therefore, epigenetic changes in the spermatozoa appear to be likely mechanisms for heritability of aspartame-induced behavioral traits. However, aspartame-induced epigenetic changes in the spermatozoa are not reported yet, although aspartame and its metabolite phenylalanine produce epigenetic changes in somatic cells ^85–87^ suggesting that aspartame has the potential to produce epigenetic changes in germ cells.

In the present study aspartame exposure did not produce changes in learned helplessness, a behavioral test employed in preclinical evaluation of antidepressant medications and considered a proxy for depression-like behavior ^88, 89^. However, whether it is a reliable measure of depression in humans is not known. Therefore, findings from studies that suggested depression-like changes following aspartame consumption in human subjects ^64, 69^ remain to be evaluated in preclinical models using appropriate behavioral measures. Similarly, aspartame exposure did not produce changes in reversal learning, memory retention or recall, despite the deficits in spatial learning and working memory. Thus, aspartame’s effects on cognitive function in our model appear to be selective to some domains but not others.

Aspartame is broken down in the gastrointestinal tract into phenylalanine, aspartic acid, and methanol. Phenylalanine crosses the blood brain barrier, and it is a precursor of monoamine neurotransmitters dopamine, epinephrine, and serotonin. These neurotransmitters regulate memory, mood, motivation, and motor function. However, previous studies reported inconsistent findings on phenylalanine and monoamine neurotransmitter levels in the blood and brain following aspartame consumption ^90–96^, suggesting that mechanisms other than monoamine neurotransmission may contribute to aspartame’s CNS effects. Alternative mechanisms such as changes in oxidative stress and gut microbiome have been proposed as well ^24, 33, 97–101^. Artificial sweeteners can alter neurotransmitter release via sweet-taste receptor activation ^102^. However, aspartame does not activate sweet taste receptors in the C57BL/6 strain of mice used here ^83, 103^. Thus, a growing list of mechanisms appear to mediate aspartame’s actions on the brain and behavior, although none has been implicated directly.

Recently, we showed that expression of genes associated with GABA and glutamate neurotransmitter signaling in the mouse amygdala is significantly influenced by daily aspartame consumption in drinking water ^35^. Specifically, genes associated with glutamatergic signaling were upregulated whereas those associated with GABA signaling were downregulated. In support of these findings, other studies showed that aspartame produced significant changes in the expression of genes associated with glutamate neurotransmission in the hypothalamus and changes in NMDA receptor signaling in the pituitary-hypothalamus-adrenal axis ^87^. Phenylalanine and aspartic acid, which are metabolites of aspartame can modulate glutamate NMDA receptor signaling ^94, 104^. Therefore, the learning and memory deficits observed in the present study may have a basis in altered glutamate and GABA neurotransmitter signaling in the brain ^35^, especially in the amygdala, which plays a key role in emotional functions, learning, and memory ^105, 106^.

The aspartame exposure paradigm used in the present study is associated with significant anxiety-like behavior in the mice ^35^. Stress and anxiety can influence the outcome from tests spatial learning and memory, which are predominantly hippocampus-based functions. Among the two well-established tests of spatial learning and memory namely the Morris water maze and Barnes maze, the outcomes from the latter test, which is used here, are less prone to the effects of stress and anxiety ^50^. Moreover, previous data showed that aspartame consumption at the dose and duration used in the present study produced significant anxiety-like behavior in the F2 generation as well, but the F2 mice in the present study did not show a robust spatial leaning deficit in the Barnes maze. If the learning and memory deficits in the Barnes maze reported here were due to the anxiety-like behavior, the deficits would have been present in the F2 generation. For these reasons we believe that anxiety associated with the aspartame exposure paradigm used here did not produce the behavioral findings in the present study.

In conclusion, aspartame consumption at doses equivalent to 7-15% of the FDA recommended maximum human daily intake value produce learning and memory deficits in male mice. These deficits are transmitted to male and female offspring in the F1 generation via the paternal lineage. Therefore, aspartame’s adverse behavioral effects may be more pervasive that currently realized, and aspartame’s safety evaluations should consider potential effects in the directly exposed individuals as well as their descendants.

## Materials and Methods

### Mouse model of aspartame exposure

C57BL/6 mice were purchased from Charles River Laboratories (Wilmington, MA) and maintained in the Florida State University Laboratory Animal Resources facility. The mice were housed in a humidity– and temperature-controlled environment on a 12-hour light-dark cycle with ad libitum access to food and water. All the protocols were approved by Florida State University Animal Care and Use Committee. All the experimental procedures and methods were carried out in accordance with our institutional guidelines and reported in accordance with ARRIVE guidelines.

Adult (approximately 8-week-old) male mice were assigned randomly to one of the following three founder (F0) groups: plain drinking water (control), 0.03% aspartame (Sigma Aldrich, catalog # A5139) containing drinking water or 0.015% aspartame in drinking water. The drinking water exposures continued for up to 16 weeks, with a fresh supply of each type of water provided weekly.

### Behavioral Analyses

Following 4, 8 and 12 weeks of exposure to the different types of drinking water, spatial working memory was evaluated using the Y-maze (Fig. 1) according to methods described previously ^35, 38, 107–110^. Briefly, the mouse was placed in one of the 3 arms of a custom-built plexiglass Y-maze with distinct visual cues at the ends of each arm. The number and sequence of arm entries over a 6-minute test period were recorded via an overhead camera. The percentage of spontaneous alternations was calculated using the following formula: [Number of correct alternations ÷ total number of arm entries] X 100. A reduction in percent spontaneous alternation reflects a spatial working memory deficit. The number of arm entries is recorded not only because it is required to calculate the percent spontaneous alternations, but also because it is a measure of general exploratory and locomotor activity. The total number of arm entries is a measure of maze performance and fewer than 6 arm entries was a criterion for exclusion. None of the mice met this criterion.

During weeks 14-15 of the exposure (Fig. 1), spatial learning and memory was assessed in the Barnes maze as described previously ^38^. Two trials per session, and one session per day were conducted using squads of 4 mice at a time. The position of the escape box remained invariant throughout the trials. Visual cues were placed prominently on all 4 walls of the testing room. Each squad contained at least one mouse from each drinking water group to control for potential variability introduced by the environment during a given trial. Each trial began with the mice placed individually at the center of the arena, covered in an upside-down metal cup, with the overhead light and fan turned on. The container was removed to release the mouse into the open arena. At the end of each trial, the maze was cleaned with a 1.6% Quatracide solution to eliminate odor cues.

During habituation, which serves to familiarize each mouse with the maze, escape box, and aversive stimuli, each mouse explored the arena in search of the escape box for 4 minutes. The mouse was gently guided to enter the escape box if it did not find it by the end of the 4 minutes. Each mouse completed one four-minute trial on each of two days of habituation.

During acquisition of learning phase, in which mice learn to locate the escape box, mice completed two 4-minute trials per day, with an intertrial interval of 15-20 minutes ^38, 39, 43^. The trial ended when the mouse entered the escape box and remained there for 30 seconds. If the mouse did not find the escape box at the end of the 4 minutes, it was gently guided into the escape box. For each trial, the time taken to locate the escape box (primary latency) was recorded. In addition, nose poke errors made during the process of finding the escape box (primary errors) were recorded. A nose poke error occurred when a mouse lowered its head and upper body into a hole that did not contain the escape box. During each trial, the strategy used by the mouse to find the escape box was also recorded. Mice use a spatial strategy by going directly to the escape hole (+/– 2 adjacent holes). The serial strategy involves visiting at least three holes adjacent to the escape box in a serial manner preceding the visit to the escape box and crossing the center of the arena one time only (i.e., the starting point). The random strategy involves unorganized search patterns including crossing the center more than one time.

Following 10 sessions (10 days) of acquisition of learning, on the 11^th^ day an acquisition probe trial was performed to evaluate spatial memory retention and retrieval. The escape box was removed, and the mouse was placed at the center of the test arena. The time spent in the quadrant that had contained the escape box in the previous 10 sessions of acquisition of learning trials (target quadrant) was recorded during the 3-minute trial. The following day, a re–acquisition of learning trial was conducted, with the same experimental setup as the previous acquisition of learning trials including return of the escape box to the original hole, to ascertain that the mice retained memory for the previously learned position of the escape box.

To analyze cognitive flexibility, the reversal learning assay was performed. The experimental setup was identical to that used during the 10 sessions of acquisition of learning, except that the escape box was moved 180° from its original position. The mice completed two, 4-minute trials per day for 4 days, with an intertrial interval of 15-20 minutes. The primary latency and primary errors were recorded. The reversal effect was calculated by comparing the difference in primary latency and primary errors between the last (10^th^) session of acquisition learning and the first session of reversal learning.

During the 16^th^ week of exposure (Fig. 1), learned helplessness was assessed using methods described previously ^111, 112^. Briefly, the mouse was suspended upside-down by its tail by taping the tail to a metal suspension rod. To prevent the mouse from climbing up its own tail, a behavior that would essentially invalidate the purpose of the test, the tail was threaded through a polycarbonate tube. The flailing movements made by the mouse to escape the stress of tail suspension and the time spent immobile (mice become immobile after several attempts to escape) were recorded over a 6-min test period. The time taken to display the first immobility event and the total time spent immobile were calculated.

The behavioral tests were performed during the active (lights-off) phase of the light-dark cycle. The data were collected and analyzed by individuals blinded to the experimental provenance of the mouse.

### Production of patrilineal F1 and F2 generations

Following 12-weeks of exposure to drinking water containing 0.015% and 0.03% aspartame, male mice from each aspartame group were bred with female mice purchased from the vendor and maintained on plain drinking water to produce the F1 generation (Fig. 1). A separate set of male mice receiving plain drinking water (control) were bred with female mice purchased from the vendor and maintained on plain drinking water to produce F1 offspring from the control, plain drinking water lineage. The F0 male mice in the aspartame and plain drinking water groups had completed three Y-maze tests (one each during weeks 4, 8 and 12) prior to breeding (Fig. 1). The 12-week duration of the aspartame exposure was chosen because our previous studies had shown that this duration of exposure was needed for heritability ^38, 70^. The female mice were checked for vaginal plugs daily and separated from the male breeder on the day the plug was discovered. Thus, female mice bred with F0 aspartame males were exposed to aspartame-containing drinking water for 1-5 days.

We bred all 8 male mice in each of the three groups (0.015% aspartame, 0.03% aspartame and plain drinking water) with female mice. The breeding resulted in 4 F1 litters from each group. We used 1-2 male and 1-2 female mice from each litter for the behavioral analyses.

To produce the F2 generation, eight F1 male mice (approximately 60-days old) each from the paternal 0.03% aspartame lineage and paternal plain drinking water lineage were bred with 2-3-months old females purchased from the vendor and maintained on plain drinking water. The F1 male mice that were bred did not perform behavioral tests but were littermates of the male F1 mice that were used for the behavioral testing. The breeding resulted in 4 F2 litters each for the 0.03% aspartame and control lineages. From each F2 litter we used 1-2 male and 1-2female mice for behavioral analyses.

F1 and F2 mice from a given drinking water lineage were housed together, and not mixed with mice from the other lineages.

### Sequence of behavioral analyses

The three Y-maze assays were performed first. Following the third Y-maze assay, F0 males were bred to produce the F1 generation. During weeks 14-15 the Barnes maze assay was performed, and during week 16, TST was performed (Fig. 1). The TST was performed last to avoid the impact of its potentially stressful experience on other behavioral assays. Following completion of the behavioral tests, the mice were anesthetized with isoflurane inhalation and sacrificed by decapitation.

### Statistical analyses

For the F0 males, we performed repeated two-way or ordinary one-way ANOVA followed by Bonferroni post hoc comparison tests, where appropriate. For the F1 and F2 generations, we used a mixed model ANOVA followed by post hoc contrasts, where appropriate, using the Benjamini-Hochberg Linear Step Up procedure to control for type I error. For the F1 and F2 generations, we treated litter as random factor (litter was a factor only for F1 and F2 but not for F0 analyses) and performed a series of mixed model ANOVA’s with drinking water exposure and sex as between subjects fixed factors. For the Barnes maze data in the F1 and F2 generations, session (day) was an additional repeated measure within subjects fixed factor. Differences among groups in the Barnes maze search strategy selection were analyzed using the nonparametric Chi-square test due to the inclusion of categorical variables. The analyses were performed using Prism 9.2.0 (GraphPad Prism, San Diego, CA) or SAS/STAT software (SAS, Carey, NC).

## Acknowledgments

This work was supported by the Jim and Betty Anne Rodgers Chair Fund.

## Author contributions

SKJ, DMM, GDS, CS and PGB designed the experiments, analyzed, and interpreted the data, and wrote the manuscript. SKJ performed the experimental work.

## Availability of Data and Materials

All data generated or analyzed during this study are included in this published article and its supplementary information files.

