## Supplementary Tables 1 and 2 for "Learning and memory deficits produced by aspartame are heritable via the paternal lineage"

**Supplementary Table 1: Litter metrics and developmental milestones**

| **Metric** | **F1 (Mean±SEM)** | | | **F2 (Mean±SEM)** | |
| --- | --- | --- | --- | --- | --- |
|  | **Control** | **0.03%** | **0.015%** | **Control** | **0.03%** |
| Litter Size | 7.60 ± 0.50 | 6.80 ± 0.40 | 7.20 ± 0.50 | 7.40 ± 0.50 | 7.00 ± 0.50 |
| P0 Weight (g) | 1.50 ± 0.05 | 1.60 ± 0.08 | 1.60 ± 0.06 | 1.70 ± 0.04 | 1.70 ± 0.06 |
| P7 Weight (g) | 3.80 ± 0.08 | 3.80 ± 0.07 | 3.90 ± 0.11 | 3.60 ± 0.10 | 3.50 ± 0.09 |
| P14 Weight (g) | 6.10 ± 0.07 | 6.20 ± 0.05 | 6.30 ± 0.07 | 6.30 ± 0.09 | 6.30 ± 0.07 |
| P21 Weight (g) | 9.50 ± 0.04 | 9.70 ± 0.06 | 9.60 ± 0.07 | 9.50 ± 0.06 | 9.50 ± 0.05 |
| External Ear Detachment (day) | 4.40 ± 0.30 | 4.10 ± 0.30 | 4.30 ± 0.30 | 4.20 ± 0.20 | 4.40 ± 0.30 |
| Fur Appearance (day) | 4.50 ± 0.30 | 4.30 ± 0.40 | 4.40 ± 0.30 | 4.10 ± 0.30 | 4.10 ± 0.30 |
| Eye Opening (day) | 13.1 ± 0.40 | 13.3 ± 0.40 | 12.9 ± 0.40 | 12.8 ± 0.40 | 13.1 ± 0.30 |

**Supplementary Table 1**: Litter size and developmental milestones were analyzed for F1 and F2 generations. The F1 generations were derived from male F0 mice from the 0.015% aspartame, 0.03% aspartame or plain drinking water (control) groups and the F2 generations were derived from 0.03% aspartame or control F1 male mice. There was no statistically significant difference between the different groups in any of the measurements (One-way ANOVA).

**Supplementary Table 2: *Post hoc* analyses**

|  | **Behavior** | **Comparison** | **F0 *t*_(df)_** | **F1 *t*_(df)_** | **F2 *t*_(df)_** |
| --- | --- | --- | --- | --- | --- |
| A | Spontaneous alternations in Y-maze | 0.015% versus 0.03% aspartame | Data presented in Table 1B | 0.41_(33)_ | Not Analyzed |
|  |  | 0.015% aspartame versus control |  | 6.24_(33)_ **** |  |
|  |  | 0.03% aspartame versus control |  | 6.64_(33)_ **** |  |
| B | Total number of arm entries in Y-maze  (*post hoc* | 0.015% versus 0.03% aspartame | Not Analyzed | 1.31_(33)_ | Not Analyzed |
|  |  | 0.015% aspartame versus control |  | 0.07_(33)_ |  |
|  |  | 0.03% aspartame versus control |  | 1.38_(33)_ |  |
| C | Barnes Maze, Acquisition of Learning: Primary errors | 0.015% Aspartame vs Water at Session 1 | 0.79_(12.78)_ | 0.99 _(171)_ | Not applicable  (0.015% aspartame lineage not available) |
|  |  | 0.015% Aspartame vs Water at Session 2 | 1.35_(11.54)_ | 4.75 _(171)_ **** |  |
|  |  | 0.015% Aspartame vs Water at Session 3 | 1.421_(13.71)_ | 5.97 _(171)_ **** |  |
|  |  | 0.015% Aspartame vs Water at Session 4 | 2.75 _(14)_* | 8.18 _(171)_ **** |  |
|  |  | 0.015% Aspartame vs Water at Session 5 | 2.41_(13.43)_ | 6.08 _(171)_ **** |  |
|  |  | 0.015% Aspartame vs Water at Session 6 | 2.83_(13.74)_* | 4.64 _(171)_ **** |  |
|  |  | 0.015% Aspartame vs Water at Session 7 | 2.65 _(10.22)_ | 3.43 _(171)_ **** |  |
|  |  | 0.015% Aspartame vs Water at Session 8 | 1.58_(12.40)_ | 2.76 _(171)_ **** |  |
|  |  | 0.015% Aspartame vs Water at Session 9 | 4.48_(10.94)_** | 3.09 _(171)_ **** |  |
|  |  | 0.015% Aspartame vs Water at Session 10 | 4.03_(12.3)_ | 2.54 _(171)_ **** |  |
|  |  | 0.03% Aspartame vs Water at Session 1 | 0.8772_(12.78)_ | 1.66 _(171)_ | 1.02 _(114)_ |
|  |  | 0.03 % Aspartame vs Water at Session 2 | 0.23_(13.92)_ | 3.21 _(171)_ **** | 0.17 _(114)_ |
|  |  | 0.03% Aspartame vs Water at Session 3 | 1.17_(11.49)_ | 6.41 _(171)_ **** | 0.51 _(114)_ |
|  |  | 0.03% Aspartame vs Water at Session 4 | 2.25_(9.65)_ | 7.63 _(171)_ **** | 0.85 _(114)_ |
|  |  | 0.03% Aspartame vs Water at Session 5 | 1.39_(13.47)_ | 5.75 _(171)_ **** | 0 _(114)_ |
|  |  | 0.03% Aspartame vs Water at Session 6 | 3.12_(13.59)_* | 4.31 _(171)_ **** | 1.19 _(114)_ |
|  |  | 0.03% Aspartame vs Water at Session 7 | 2.82_(9.98)_ | 3.43 _(171)_ **** | 0.51 _(114)_ |
|  |  | 0.03% Aspartame vs Water at Session 8 | 1.29_(10.79)_ | 3.65 _(171)_ **** | 0.17 _(114)_ |
|  |  | 00.03% Aspartame vs Water at Session 9 | 4.18_(12.31)_** | 3.32 _(171)_ **** | 0.68 _(114)_ |
|  |  | 0.03% Aspartame vs Water at Session 10 | 4.15_(11.12)_ | 2.43 _(171)_ **** | 0.51 _(114)_ |
|  |  | 0.015% vs 0.03% aspartame at Session 1 | 0.26_(11.77)_ | 0.66 _(171)_ | Not applicable  (0.015% aspartame lineage not available) |
|  |  | 0.015% vs 0.03% aspartame at Session 2 | 1.14_(12.07)_ | 1.55 _(171)_ |  |
|  |  | 0.015% vs 0.03% aspartame at Session 3 | 0.49_(12.5)_ | 0.44 _(171)_ |  |
|  |  | 0.015% vs 0.03% aspartame at Session 4 | 1.31_(9.61)_ | 0.55 _(171)_ |  |
|  |  | 0.015% vs 0.03% aspartame at Session 5 | 1.14_(14)_ | 0.33 _(171)_ |  |
|  |  | 0.015% vs 0.03% aspartame at Session 6 | 0.75_(12.81)_ | 0.33 _(171)_ |  |
|  |  | 0.015% vs 0.03% aspartame at Session 7 | 0.22_(13.97)_ | 0 _(171)_ |  |
|  |  | 0.015% vs 0.03% aspartame at Session 8 | 0.52_(13.29)_ | 0.88 _(171)_ |  |
|  |  | 0.015% vs 0.03% aspartame at Session 9 | 0.77_(13.48)_ | 0.22 _(171)_ |  |
|  |  | 0.015% vs 0.03% aspartame at Session 10 | 0.52_(13.61)_ | 0.11 _(171)_ |  |
| D | Barnes Maze, Acquisition of Learning: Primary latency | 0.015% Aspartame vs Water at Session 1 | 0.86_(10.51)_ | 1.27 _(171)_ | Not applicable  (0.015% aspartame lineage not available) |
|  |  | 0.015% Aspartame vs Water at Session 2 | 1.13_(10.3)_ | 6.29 _(171)_ **** |  |
|  |  | 0.015% Aspartame vs Water at Session 3 | 2.93_(12.9)_* | 7.63 _(171)_ **** |  |
|  |  | 0.015% Aspartame vs Water at Session 4 | 1.89_(13.38)_ | 7.49 _(171)_ **** |  |
|  |  | 0.015% Aspartame vs Water at Session 5 | 2.82_(13.99)_* | 5.32 _(171)_ **** |  |
|  |  | 0.015% Aspartame vs Water at Session 6 | 2.13(_13.86)_ | 5.9 _(171)_ **** |  |
|  |  | 0.015% Aspartame vs Water at Session 7 | 3.60_(13.99)_** | 4.23 _(171)_ **** |  |
|  |  | 0.015% Aspartame vs Water at Session 8 | 3.90_(10.37)_ | 2.72 _(171)_ **** |  |
|  |  | 0.015% Aspartame vs Water at Session 9 | 1.90_(9.60)_** | 3.26 _(171)_ **** |  |
|  |  | 0.015% Aspartame vs Water at Session 10 | 2.48_(11.96)_ | 1.6 _(171)_ |  |
|  |  | 0.03% Aspartame vs Water at Session 1 | 0.86_(10.51)_ | 1.64 _(171)_ | 0.53 _(114)_ |
|  |  | 0.03 % Aspartame vs Water at Session 2 | 0.09_(13.86)_ | 5.8 _(171)_ **** | 1.16 _(114)_ |
|  |  | 0.03% Aspartame vs Water at Session 3 | 1.86_(13.38)_ | 7.65 _(171)_ **** | 1.41 _(114)_ |
|  |  | 0.03% Aspartame vs Water at Session 4 | 1.08_(12.38)_ | 8.44 _(171)_ **** | 0.55 _(114)_ |
|  |  | 0.03% Aspartame vs Water at Session 5 | 1.56_(13.97)_ | 5.49 _(171)_ **** | 0.56 _(114)_ |
|  |  | 0.03% Aspartame vs Water at Session 6 | 3.77_(11.68)_** | 5.8 _(171)_ **** | 0.79 _(114)_ |
|  |  | 0.03% Aspartame vs Water at Session 7 | 4.58_(13.65)_** | 4.08 _(171)_ **** | 0.25 _(114)_ |
|  |  | 0.03% Aspartame vs Water at Session 8 | 2.16_(13.56)_ | 3.51 _(171)_ **** | 0.32 _(114)_ |
|  |  | 00.03% Aspartame vs Water at Session 9 | 1.67_(9.42)_ | 3.29 _(171)_ **** | 0.79 _(114)_ |
|  |  | 0.03% Aspartame vs Water at Session 10 | 2.44_(10.9)_ | 1.09 _(171)_ | 0.05 _(114)_ |
|  |  | 0.015% vs 0.03% aspartame at Session 1 | 0.17_(11.37)_ | 0.37 _(171)_ | Not applicable  (0.015% aspartame lineage not available) |
|  |  | 0.015% vs 0.03% aspartame at Session 2 | 0.93_(9.74)_ | 0.5 _(171)_ |  |
|  |  | 0.015% vs 0.03% aspartame at Session 3 | 0.47_(11.40)_ | 0.02 _(171)_ |  |
|  |  | 0.015% vs 0.03% aspartame at Session 4 | 1.07_(13.66)_ | 0.95 _(171)_ |  |
|  |  | 0.015% vs 0.03% aspartame at Session 5 | 1.33_(13.94)_ | 0.17 _(171)_ |  |
|  |  | 0.015% vs 0.03% aspartame at Session 6 | 0.99_(10.39)_ | 0.10 _(171)_ |  |
|  |  | 0.015% vs 0.03% aspartame at Session 7 | 0.77_(13.77)_ | 0.14 _(171)_ |  |
|  |  | 0.015% vs 0.03% aspartame at Session 8 | 1.59_(11.55)_ | 0.79 _(171)_ |  |
|  |  | 0.015% vs 0.03% aspartame at Session 9 | 0.14_(13.98)_ | 0.03 _(171)_ |  |
|  |  | 0.015% vs 0.03% aspartame at Session 10 | 0.17_(13.68)_ | 0.51 _(171)_ |  |
| E | Barnes Maze, Reversal Learning - 4 sessions: Primary errors | 0.015% Aspartame vs Water at Session 1 | 1.47_(9.96)_ | 0.97 _(63)_ | N/A |
|  |  | 0.015% Aspartame vs Water at Session 2 | 1.94_(8.89)_ | 1.08 _(63)_ |  |
|  |  | 0.015% Aspartame vs Water at Session 3 | 3.23_(13.91)_* | 1.51 _(63)_ |  |
|  |  | 0.015% Aspartame vs Water at Session 4 | 1.71_(12.31)_ | 0.11 _(63)_ |  |
|  |  | 0.03% Aspartame vs Water at Session 1 | 2.16_(9.33)_ | 1.99 _(63)_ | 0.32 _(30)_ |
|  |  | 0.03 % Aspartame vs Water at Session 2 | 2.56_(8.77)_ | 2.369 _(63)_** | 0.21 _(30)_ |
|  |  | 0.03% Aspartame vs Water at Session 3 | 2.87_(14.0)_* | 0.86_(63)_ | 0.42 _(30)_ |
|  |  | 0.03% Aspartame vs Water at Session 4 | 2.68_(13.51)_ | 0.32 _(63)_ | Session 4 not analyzed |
|  |  | 0.015% vs 0.03% aspartame at Session 1 | 0.68_(13.77)_ | 1.03 _(63)_ | Not applicable  (0.015% aspartame lineage not available) |
|  |  | 0.015% vs 0.03% aspartame at Session 2 | 0.51_(13.98)_ | 1.62 _(63)_ |  |
|  |  | 0.015% vs 0.03% aspartame at Session 3 | 0.31_(13.87)_ | 0.65 _(63)_ |  |
|  |  | 0.015% vs 0.03% aspartame at Session 4 | 0.61_(13.49)_ | 0.22 _(63)_ |  |
| F | Barnes Maze, Reversal Learning - 4 sessions: Primary latency | 0.015% Aspartame vs Water at Session 1 | 1.02_(13.38)_ | 0.37 _(63)_ | Not applicable  (0.015% aspartame lineage not available) |
|  |  | 0.015% Aspartame vs Water at Session 2 | 1.64_(13.99)_ | 0.84 _(63)_ |  |
|  |  | 0.015% Aspartame vs Water at Session 3 | 2.10_(13.17)_ | 0.04 _(63)_ |  |
|  |  | 0.015% Aspartame vs Water at Session 4 | 2.44_(13.93)_ | 0.2 _(63)_ |  |
|  |  | 0.03% Aspartame vs Water at Session 1 | 0.38_(13.36)_ | 0.99 _(63)_ | 0.17 _(30)_ |
|  |  | 0.03 % Aspartame vs Water at Session 2 | 2.06_(10.48)_ | 0.69 _(63_ | 1.69 _(30)_ |
|  |  | 0.03% Aspartame vs Water at Session 3 | 2.37_(13.86)_ | 0.07_(0.0763)_ | 0.69 _(30)_ |
|  |  | 0.03% Aspartame vs Water at Session 4 | 1.50_(12.15)_ | 0.04 _(63)_ | Session 4 not analyzed |
|  |  | 0.015% vs 0.03% aspartame at Session 1 | 0.47_(11.94)_ | 0.62 _(63)_ | Not applicable  (0.015% aspartame lineage not available) |
|  |  | 0.015% vs 0.03% aspartame at Session 2 | 0.02_(10.61)_ | 0.14 _(63)_ |  |
|  |  | 0.015% vs 0.03% aspartame at Session 3 | 0.07_(13.68)_ | 0.12 _(63)_ |  |
|  |  | 0.015% vs 0.03% aspartame at Session 4 | 0.35_(11.67)_ | 0.16 _(63)_ |  |
| G | Total time immobile in Tail Suspension Test | 0.015% versus 0.03% aspartame | N/A | 0.04 _(33)_ | Test not performed |
|  |  | 0.015% aspartame versus control |  | 0.62 _(33)_ |  |
|  |  | 0.03% aspartame versus control |  | 0.66 _(33)_ |  |

**Supplementary Table 2**: *Post hoc* contrast analysis of the data from Y-maze (spatial working memory), Barnes maze (spatial learning and memory) and tail suspension test (learned helplessness) between the aspartame and plain drinking water (control) lineages in the F1 and F2 generations. Bonferroni multiple comparisons test was used for the F0 data whereas the Benjamini-Hochberg Linear Step Up procedure was used to control for type I error for the F1 and F2 data.
